# Goal-directed action selection relies on conjunctive representations that bind goals, actions and expected sensory outcomes

**DOI:** 10.64898/2026.07.24.740508

**Authors:** Jet Lageman, Atsushi Kikumoto, David Badre, Johannes J. Fahrenfort, Heleen A. Slagter

## Abstract

Goal-directed actions are performed to obtain specific sensory outcomes. However, theories of action control differ in how the brain encodes action-outcome predictions and whether they are encoded as separable components or integrated into conjunctive representations that support behavior and learning. Here, we tested whether predicted sensory outcomes are integrated into the neural representations underlying the selection of goal-directed actions. Participants learned novel action-outcome contingencies and selected actions to generate specific sensory outcomes, while their brain activity was recorded using EEG. Combining time-resolved multivariate decoding with representational similarity analysis, we tracked both feature-specific and conjunctive neural representations over time. We found that expected sensory outcomes were integrated into high-dimensional conjunctive representations that could not be explained by a linear addition of individual features. The strength of these representations predicted trial-to-trial reaction times, alongside compositional action representations. Over time, the strength of compositional feature representations weakened, suggesting that the brain more selectively relied on conjunctive representations over the course of learning. These results provide neural evidence that predicted sensory outcomes are incorporated into behaviorally relevant conjunctive representations during action selection, consistent with theories that propose integrated action-outcome representations.

## Introduction

Goal-directed action is commonly defined as grounded in knowledge about action-outcome contingencies (*1*). However, there is a gap in our understanding of how the brain represents such action-outcome relationships and how these representations contribute to flexible action selection. Traditional motor command theories propose that sensory outcomes are predicted by an internal forward model that transforms planned motor commands into expected outcomes (*2–9*), used to overcome sensorimotor delays and reduce uncertainty in state estimates (*2*, *5*, *9*). Ideomotor and related action–effect theories ascribe a more fundamental role to sensory outcome predictions, proposing that actions are selected in terms of their anticipated sensory consequences, with bidirectional associations linking actions and sensory outcomes (*10–14*). Such theories suggest that the brain may integrate goals, actions, and expected sensory outcomes into high-dimensional conjunctive representations (*15*) or ‘event files’ (*13*, *14*, *16*). These event files are proposed to integrate relevant features (e.g., abstract goals, sensory information, actions, outcomes) into a common representational format, whereby the activation of one feature triggers the activation of all bound features (*12*, *17–20*), allowing the brain to flexibly learn different action-outcome contingencies. Active inference offers a more formal computational framework for this idea by conceptualizing action selection as the control of sensory input (*21*, *22*). Within this framework, actions are specified in terms of their expected sensory consequences (*23–27*) and implemented through cranial nerve and spinal reflex mechanisms (*28*, *29*), such that behavior minimizes the mismatch between predicted and actual sensory input. This implies a tight integration of anticipated sensory outcomes and action plans during action selection.

Despite the longstanding consensus that outcome predictions contribute to action control, it remains unclear how the brain encodes such predictions. Prior work on action-outcome predictions has shown that expected sensory consequences of actions can be decoded from neural activity prior to action onset (*26*, *30–36*). However, the representational format of these predictions remains unclear. Specifically, it is unknown whether anticipated outcomes are represented primarily through the interaction of separable, compositional codes for goals, actions, and outcomes, or through integrated conjunctive representations that bind these features together into event-file-like structures. Answering this question requires the investigation of neural representations during action control, both in terms of content (i.e., *what* is represented) and geometry (i.e., *how* is something represented) (*37–39*).

One possibility is that action-outcome relationships are encoded in a conjunctive format that combines multiple features, its high embedding dimensionality permitting flexible separation of multiple task-specific action-outcome predictions that vary over time and context (*38*, *40–42*). However, a downside of high-dimensional conjunctive codes or event files is that they are more sensitive to small input variations and slower to generalize to novel contexts, unlike more low-dimensional compositional codes that code for a single task-relevant feature while abstracting over other features (*43–47*). Thus, another possibility is that actions are represented in a low-dimensional and compositional format (*48–50*), resembling modular ‘motor primitives’ (*51–53*), that may be flexibly combined with higher-level context representations and goals and lower-level outcome predictions, without requiring integration of their neural codes.

Recent EEG studies have identified a potential neural marker of event files during tasks that required flexible switching between stimulus-response rules (*15*, *54*, *55*). Using a novel analysis technique combining linear decoding with representational similarity analysis (RSA) (*39*, *56*), these studies showed that several task-relevant features, including abstract rules, stimuli, and responses, were non-linearly integrated into conjunctive representations. The trial-to-trial strength of these conjunctive representations predicted efficient action control, including action selection (*54*), inhibition (*57*), and flexible task learning (*58*, *59*). However, it is unknown whether similar conjunctive representations can also incorporate predicted action outcomes to support goal-directed action selection.

The aim of the current EEG study was to determine how the brain represents sensory outcome predictions and whether goal-directed action selection fundamentally incorporates these predictions, by examining if action-outcome learning is associated with the development of high-dimensional, outcome-conjunctive neural representations that predict behavior. While their brain activity was recorded with EEG, participants performed goal-directed actions to generate specific sensory outcomes (Fig 1A). Goal-directed action-outcome relationships pose a challenge for identifying conjunctive representations, as goals tend to overlap with expected sensory outcomes. For example, when pressing a doorbell, the bell sound is both the goal and predicted outcome. Therefore, we designed our task so that we could differentiate predicted sensory outcomes from goals, and determine whether action selection is supported by representations that incorporate outcome features associated with a planned action, but irrelevant to the current goal. If such representations contribute to behavior, this would provide strong evidence that the brain encodes integrated action-outcome predictions rather than merely representing the intended goal. Combining linear decoding with representational similarity analysis, we tracked the representational strength of individual task features over time, as well as their non-linear integration into high-dimensional conjunctive representations (*15*, *38*), to determine whether goals, actions, and predicted outcomes are mostly coded in an abstract, compositional format or in a context-sensitive, conjunctive manner. Moreover, we examined whether action-outcome integration contributes to action selection. If this is the case, one may predict that the integration of actions and sensory outcomes has functional relevance for action selection, and does not merely emerge as the product of a separate state estimation module. Our results show that outcome predictions are encoded in high-dimensional, context-sensitive representations, and that the integration of anticipated sensory outcomes into conjunctive representations robustly predicts performance in goal-directed action selection over the course of learning, suggesting a critical role for action-outcome predictions in goal-directed action selection.

**Fig 1.**
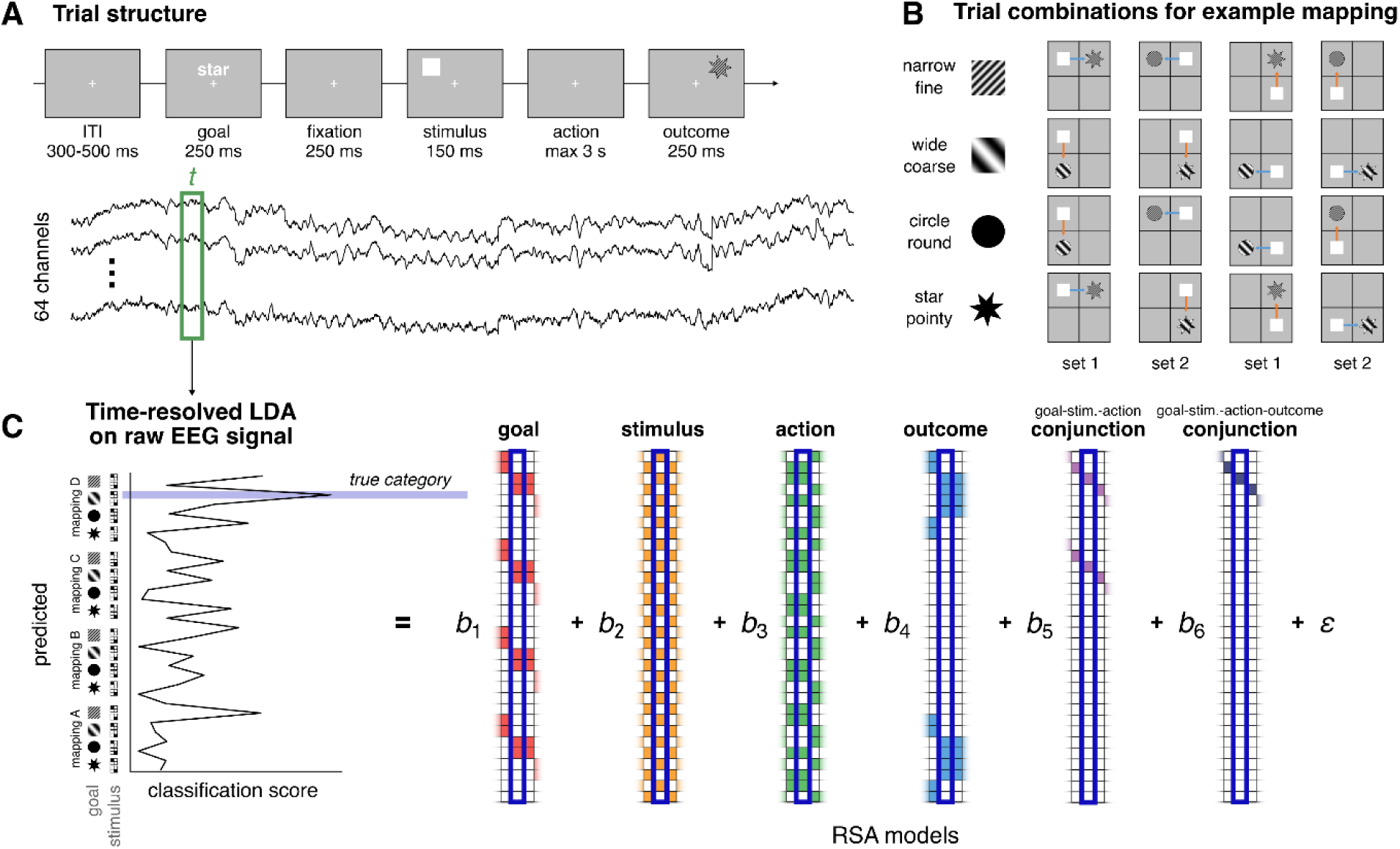
Task and analysis structure. **(A)** Sequence of events in the action-outcome learning task. After a goal cue (e.g., *star*), a stimulus appeared on one of four starting locations. Participants then selected an action to move the stimulus either vertically or horizontally by pressing one of two buttons, to generate an outcome at the target location (e.g., *narrow star*). **(B)** All possible trial combinations for a single example location-outcome mapping, illustrating how stimuli can be moved to generate the correct outcome under each goal. Two distinct cues were used for each goal. For decoding analyses, trial combinations were divided into two independent sets: one containing all trials with stimuli in the upper-left or lower-right locations and another containing all trials with stimuli in the upper-right or lower-left locations. **(C)** Schematic overview of the decoding and representational similarity analysis. For each trial-specific time point *t*, a classifier was trained on raw EEG data from 64 channels to distinguish specific trial combinations, yielding vectors of 32 classification scores. These vectors were then regressed onto the vectors from 6 RSA model matrices, each representing a possible underlying similarity structure (see Fig S1), and 3 control vectors. The resulting single-trial *t*-values quantify how strongly each model explains the observed pattern of similarities over time.

## Results

40 participants performed an action-outcome learning task (Fig 1A) across two sessions on separate days. This sample size was based on previous work, in which robust results were found for sample sizes of approximately 24 participants (*15*, *54*, *57*), although we increased the sample size and the number of experimental trials due to the higher number of possible conjunctions in the current study. Each session comprised 12 blocks of at least 160 trials, with a hidden location-outcome mapping that changed every block. On every trial, participants were asked to generate an outcome with a specified goal feature (e.g., *star*) by moving a stimulus either horizontally or vertically to a new location using one of two button presses. The resulting location determined the identity of the outcome generated under the current mapping (e.g., *narrow star*). As illustrated in Fig 1B, stimuli in each location can be acted upon to reach every goal under a given location-outcome mapping. Notably, the same outcome could be produced from different starting locations and under different goals (e.g., a narrow star could be generated when the goal was star *or* narrow). This design allowed us to dissociate between neural representations related to goals and sensory outcomes: the goal-representation should only track only the *goal-relevant* dimension of the outcome (e.g., shape, when the goal is *star*) whereas the outcome representation should track both dimensions. Moreover, the disassociation of goals and sensory outcomes enabled us to separately track conjunctive representations integrating goal-stimulus-action combinations (i.e., only including the goal-relevant dimension) from those additionally integrating predicted sensory outcomes (i.e., integrating both goal-relevant and goal-irrelevant dimensions of the sensory outcome). In addition, the use of multiple location-outcome mappings allowed us to disentangle the location of a particular outcome from its identity. Because a goal-directed action specifies not only what outcome is desired, but also where it is expected to occur, we explicitly controlled for outcome location in our analysis. This allowed us to isolate representations of predicted outcome identity, which in our design was independent from the stimulus, action, and specific stimulus-action combinations, from representations of the spatial target of the action and determine whether sensory outcome information contributes to action selection beyond anticipation of outcome location. Critically, this way, we could determine whether action selection depends only on representations of goal-relevant outcome information, or whether it also incorporates goal-irrelevant sensory outcome features that are associated with the selected action. Evidence for the latter would provide strong support for genuinely outcome-integrating event representations, rather than representations that simply encode the goal-relevant feature or anticipated outcome location.

### Behavior

Fig 2 shows that participants quickly learned how to generate goal outcomes, with performance stabilizing within a few blocks. Both accuracy and reaction times (RTs) were best explained by a linear model with main and interaction effects that included learning at three levels: across sessions, across blocks within sessions, and across trials within blocks (ACCURACY: *session: b* = 0.218, *t*_161,901_ = 26.13, *p* < 0.001, *block: b* = 0.076, *t*_161,901_ = 9.26, *p* < 0.001, *trial: b* = 0.049, *t*_161,901_ = 6.30, *p* < 0.001; *session × block: b* = -0.059, *t*_161,901_ = -7.077, *p* < 0.001; *session × trial: b* = 0.006, *t*_161,901_ = 0.724, *p* = 0.469; *block × trial: b* = -0.050, *t*_161,901_ = -5.983, *p* < 0.001; *session × block × trial: b* = 0.019, *t*_161,901_ = 2.256, *p* = 0.024; REACTION TIME *session: b* = -0.126, *t*_161,901_ = -121.21, *p* < 0.001, *block: b* = -0.048, *t*_161,901_ = -46.61, *p* < 0.001, *trial: b* = -0.020, *t*_161,901_ = -18.61, *p* < 0.001; *session × block: b* = 0.033, *t*_161,901_ = 31.950, *p* < 0.001; *session × trial: b* = 0.010, *t*_161,901_ = 9.645, *p* < 0.001; *block × trial: b* = 0.008, *t*_161,901_ = 7.147, *p* < 0.001; *session × block × trial: b* = -0.0004, *t*_161,901_ = -0.377, *p* = 0.706). The most substantial behavioral improvements seem to take place during the first blocks of the first session, as supported by the significant interaction effects between session × block and between block × trial, and due to offline gain between the two sessions, as supported by the significant main effects of session number on accuracy and RT.

**Fig 2.**
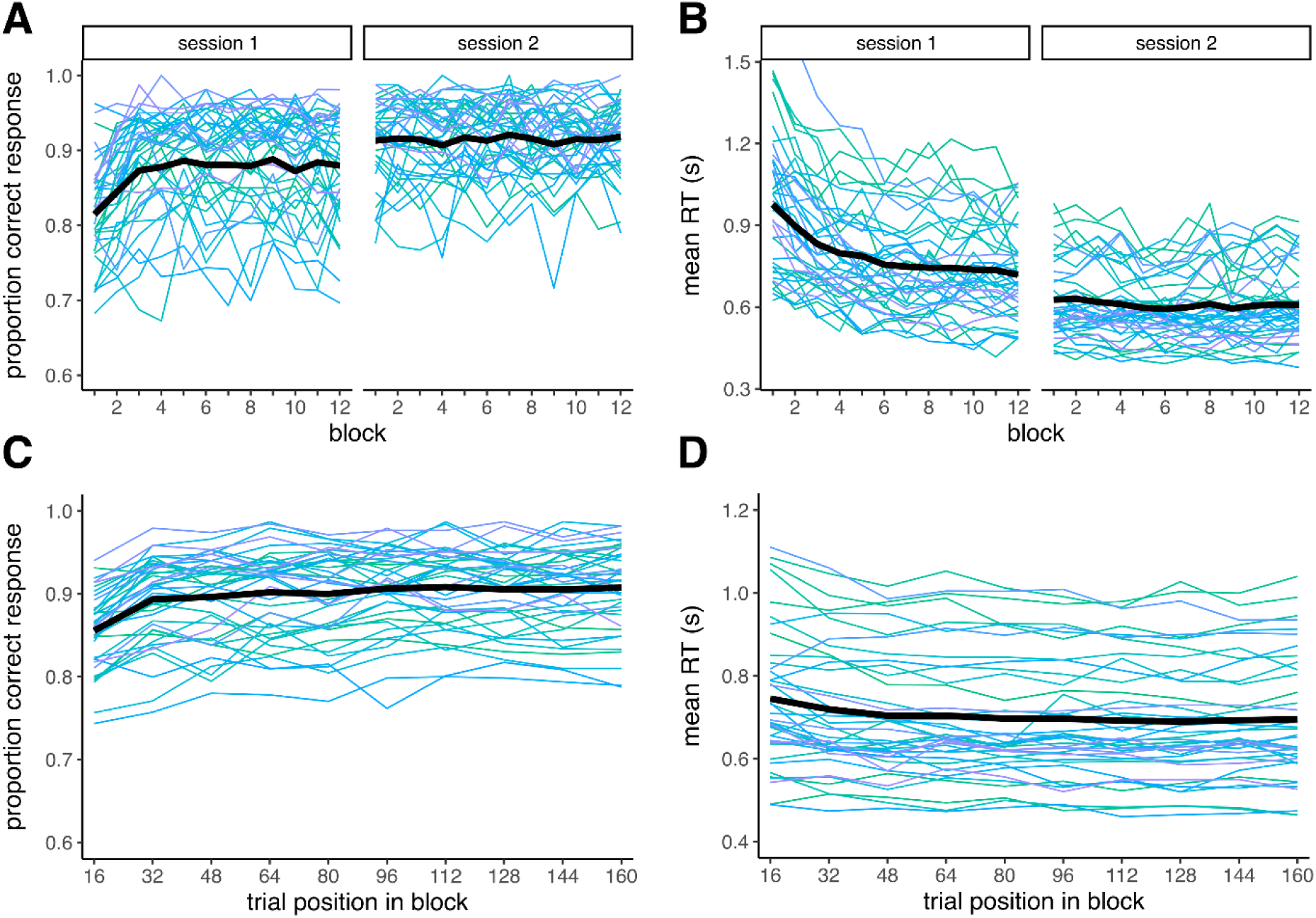
Behavioral results. **(A)** Accuracy (proportion correct response) across blocks and sessions. **(B)** Mean reaction times in seconds across blocks and sessions. **(C)** Accuracy (proportion correct response) as a function of trial position within block. **(D)** Mean reaction times in seconds as a function of trial position within block. Black lines represent grand average, colored lines represent individual participants.

### Representational dynamics

To test whether action selection relies on the integration of sensory outcome predictions into conjunctive neural representations, we combined multivariate pattern analysis (MVPA) and representational similarity analysis (RSA) (*56*). This approach allowed us to characterize the representational structure of neural activity in a quantitative manner with RSA, teasing apart contributions from correlated sources, while using the high temporal resolution of EEG-based MVPA to track when specific feature representations emerged. We quantified the activation of neural representations reflecting the current trial goal, stimulus location, required action, predicted outcome identity, and two types of conjunctive representations: those integrating goal-stimulus-action features and those additionally integrating predicted sensory outcomes (i.e., goal-stimulus-action-outcome relationships) (Fig S1A). To ensure that observed similarities were not driven by design constraints, we included additional RSA model matrices capturing location-outcome mappings (to control for the fact that unique trial combinations would only occur in specific mappings), and outcome locations (to control for neural patterns that were predictive of a particular outcome location rather than its identity). Moreover, we included the subject-specific mean RT for each unique trial combination as a nuisance regressor to account for idiosyncratic differences between unique trial combinations unrelated to the composition of task features (Fig S1B).

Fig 3A shows that decoded features at the level of single trials unfolded in line with the temporal structure of the task and in line with single-feature classifier results (Fig S2), with a range of *t*-values comparable to those found in previous studies that employed the same analysis (*15*, *54*, *57*, *60*). Note that *t*-values here should be interpreted not in the usual way, but as the relative contributions of a given predictor RSA matrix. In the stimulus-locked analysis, a goal-related representation emerged shortly after the goal cue, followed by strong decoding of the stimulus location after stimulus onset, and subsequent emergence of action-related and conjunctive goal-stimulus-action representations. In the action-locked analysis, stimulus location representations again dominate early on, followed by peaks of activation for the action and goal-stimulus-action representations in the period before action onset. In neither case did we observe significant activation of outcome identity representations or goal-stimulus-action-outcome representations, despite being able to decode outcome identities using MVPA (Fig S2). However, goal-stimulus-action-outcome representations were present in individual trials, showing a range of *t*-values similar to single feature representations. This suggests that the absence of a sustained increase in the average time course may be explained by larger variability of activations across time samples or across trials.

**Fig 3.**
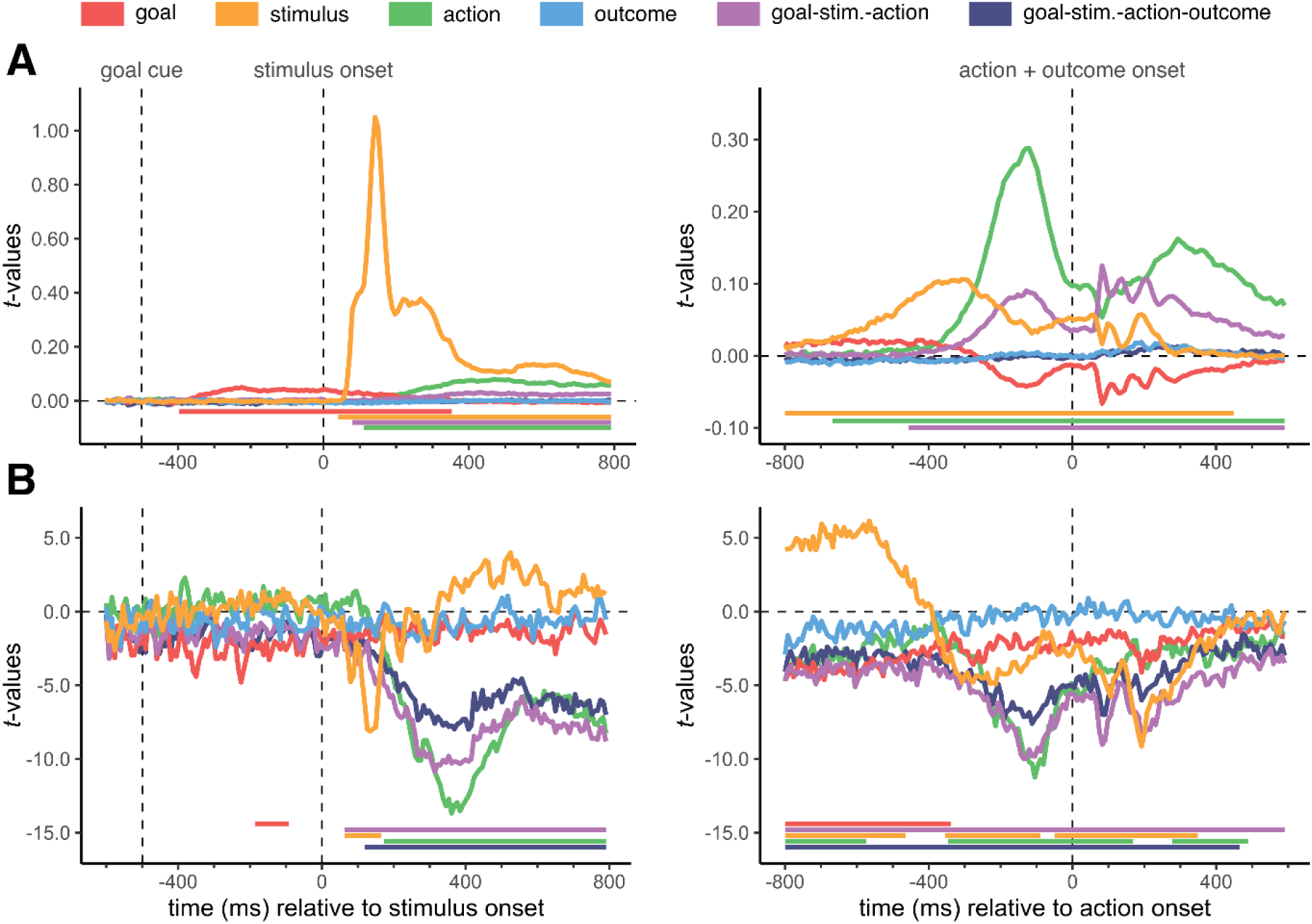
Representational time course. **(A)** Average trajectories of single-trial *t*-values from the representational similarity analysis, time-locked to stimulus onset (*left*) and action onset (*right*). Higher *t*-values indicate that a given feature (e.g., stimulus location) or combination of features explains more variance in the EEG similarity structure, when controlling for all other features. Note that the range of the Y-axis differs between the left and right subplot. **(B)** Trajectories of *t*-values from mixed linear models predicting trial-to-trial RTs from RSA-derived feature scores, time-locked to stimulus onset (*left*) and action onset (*right*). More negative *t*-values indicate that stronger feature-specific or conjunctive RSA patterns are associated with faster trial-to-trial RTs, controlling for all other predictors. Colored bars beneath each panel denote significant clusters identified via nonparametric cluster-based permutation testing (1000 permutations; cluster-forming threshold: *p* < 0.05; cluster significance threshold: *p* < 0.01).

However, we found that conjunctive representations including sensory outcome predictions predicted behavior in a next analysis examining how the time-resolved activation strength of feature-specific and conjunctive neural representations (expressed in *t*-values) related to trial-to-trial RTs. As shown in Fig 3B, stronger action, goal-stimulus-action, and goal-stimulus-action-outcome representations predict faster RTs, with the clearest effects emerging between 300-400 ms after stimulus presentation or around 100 ms before action onset. Crucially, conjunctive representations that included predicted sensory outcomes explained variance in behavioral performance, even when controlling for the effects of individual task features such as the outcome location and the goal-stimulus-action conjunction. Moreover, the timing of these effects, peaking before action execution, suggest they reflect processes related to action preparation rather than separate, downstream perceptual consequences. The simultaneous effects of action representations and conjunctive representations could indicate that the brain concurrently maintains feature representations in compositional and conjunctive formats (*61–63*). Importantly, however, we only observed representations of outcome identity in a conjunctive format, and with a functional relationship to behavior, consistent with integrative theories of action selection.

Although outcome location effects were explicitly controlled for in the RSA analysis and implicitly in the experimental design, in which stimulus and target locations were combined with different goals and outcomes, the peripheral presentation of stimuli raises the possibility that part of the EEG signal reflects eye movements rather than brain activity. To more definitively rule this out, we repeated our MVPA analysis using the 4 recorded EOG channels and 3 eye-tracking measures (X-coordinate of gaze position, Y-coordinate of gaze position, pupil size). As shown in Fig S3, this control analysis showed that stimulus location could be reliably decoded from eye channels, but subsequent regression analyses did not reproduce the relationships between decoded feature representations and trial-to-trial RTs observed in the EEG-based analysis. This indicates that eye movements cannot account for our main findings.

### Learning effects

We next assessed whether representational dynamics changed as participants improved their performance on the task both over the course of the two sessions, and within blocks, as they learned each block’s action-outcome mapping. To this end, we examined whether the strength of feature- and conjunction-specific representations (expressed as *t*-values) varied systematically as a function of trial position, both across blocks and sessions and within a single block. The resulting time-courses are depicted in Fig 4.

**Fig 4.**
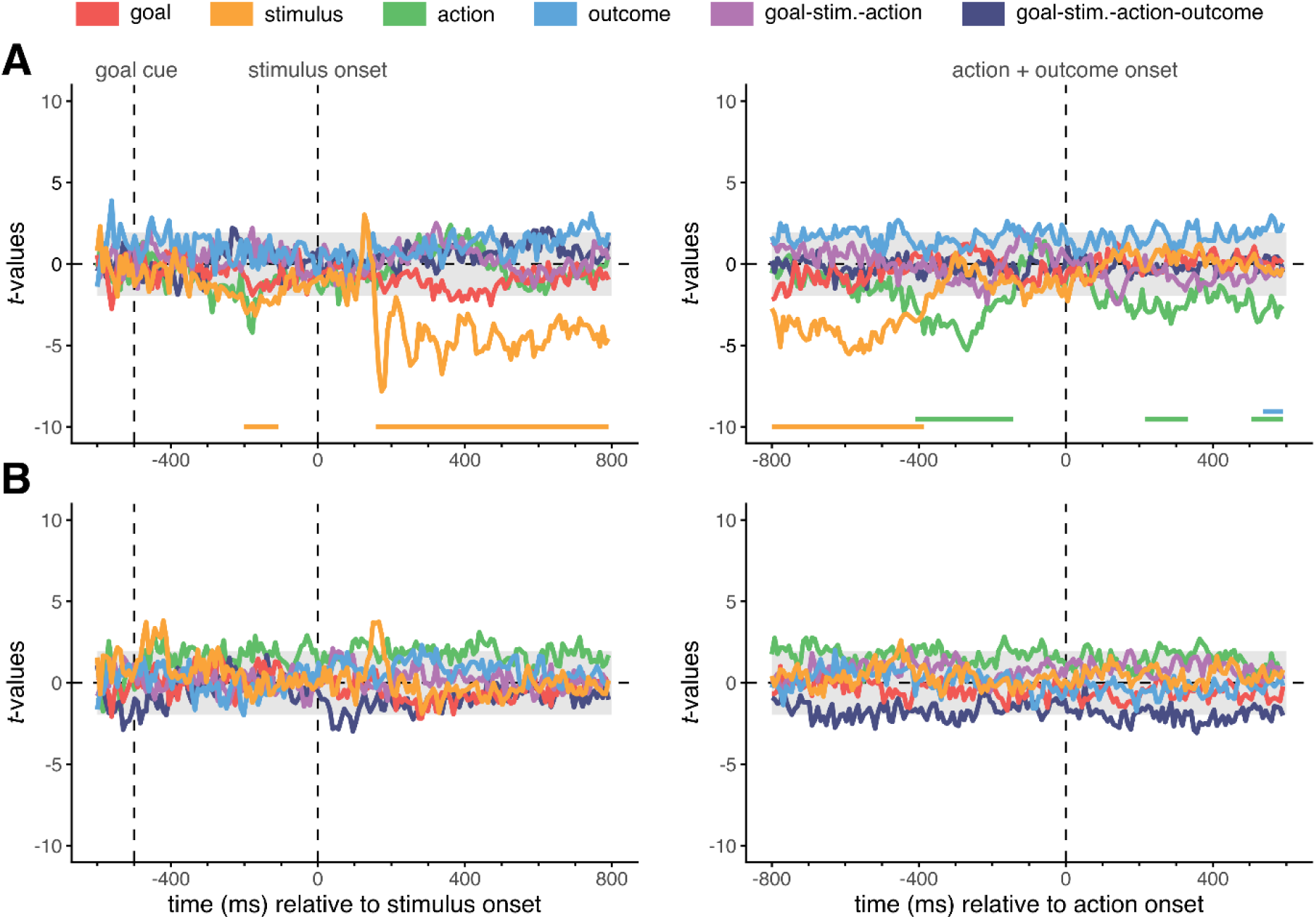
Development of representations over trials. **(A)** Trajectories of *t*-values from linear models predicting feature-specific or conjunctive representational strength as a function of trial position *across* blocks and sessions, time-locked to stimulus onset (*left*) and action onset (*right*). **(B)** Trajectories of *t*-values from mixed linear models predicting feature-specific or conjunctive representational strength as a function of trial position *within* a block, time-locked to stimulus onset (*left*) and action onset (*right*). Shaded gray areas represent the cluster-forming threshold of *p* < 0.05; colored bars beneath each panel denote significant clusters identified via nonparametric cluster-based permutation testing (1000 permutations; cluster significance threshold: *p* < 0.01).

Across all trials in the experiment (Fig 4A), the most striking changes are a decrease in the stimulus representation strength from about 150 ms after stimulus onset throughout the stimulus-locked epoch and a decrease in the action representation strength from about 400 ms to 150 ms before action onset. The decreasing strength of these low-dimensional representations could indicate a shift from compositional to conjunctive coding over the course of learning, in line with past reports for stimulus-response selection (*42*, *60*, *64*), although we do not see a corresponding increase in the strength of conjunctive representations. The stability of these conjunctive representations over time suggests that trial-to-trial variability, not gradual representational change over the course of learning, drives the observed link between conjunctive representation strength and performance, both for goal-stimulus-action and goal-stimulus-action-outcome conjunctions.

For learning within blocks (Fig 4B), nonparametric cluster-based permutation testing did not yield any significant clusters, suggesting that representational dynamics were relatively stable over the course of single blocks. This result mirrors the limited behavioral improvements within single blocks, relative to the larger improvements within blocks and sessions. It may also indicate that the brain’s representational geometry adapts to task demands only over longer timescales, in line with previous findings (*60*).

## Discussion

How does the brain predict sensory outcomes during goal-directed action selection? Here, by combining time-resolved multivariate decoding with representational similarity analysis, we show that high-dimensional conjunctive representations integrating goals, stimuli, actions, and their anticipated sensory outcomes can be identified at a single-trial level before action execution. The moment-to-moment strength of such representations predicted trial-to-trial reaction times, even after accounting for individual task features and for conjunctive patterns without outcome information. Together, these results provide neural evidence for the notion that event files, in the form of high-dimensional integrated representations containing context-sensitive sensory outcome predictions, actively support the selection of goal-directed actions. Our findings emphasize that predicted sensory outcomes are not downstream consequences, but core determinants of action selection, consistent with integrative accounts such as active inference in which action planning concerns the control of expected sensory input.

Our results extend previous work showing that conjunctive representations of context-dependent stimulus-response associations support flexible action selection (*15*, *54*, *55*, *65*), by demonstrating that such high-dimensional conjunctive representations can also incorporate predicted sensory action outcomes. This result is striking, because the outcome dimension we decode is not strictly task-relevant and cannot be reduced to the goal feature itself. In our design, the goal-relevant feature of predicted sensory outcomes (e.g., shape, when the goal is ‘star’) is already accounted for by both the goal representation and the conjunctive goal-stimulus-action representation. Moreover, effects specific to the location of the outcome rather than its identity have been regressed out. Therefore, the additional explanatory power of the goal-stimulus-action-outcome conjunction is fully attributable to the integration of the goal-irrelevant outcome feature (e.g., the spatial frequency of the ‘star’ outcome). This shows that the brain binds together predicted sensory consequences even when those consequences do not constrain the required action and do not define the goal, a hallmark of genuinely outcome-integrating event structures rather than simple goal coding. Moreover, by including nuisance regressors for idiosyncrasies in RT patterns of unique conjunctions, specific location-outcome mappings, and outcome locations, and by demonstrating that eye movement signals do not reproduce these effects, we rule out alternative explanations, such as statistical regularities or peripheral artifacts, for the relationship between the conjunctive representations and performance. Together, these results provide strong evidence that conjunctive representations supporting goal-directed action selection incorporate predicted sensory outcomes in a functionally significant manner.

Motor command theories and integrative theories of action selection both account for flexible, context-sensitive behavior, but these lines of research have traditionally suggested different neural and representational architectures supporting this behavior. Research following motor command theories has more commonly described the representational structure of action control as hierarchical and compositional (*3*, *48*, *49*, *66*), with abstract representations of tasks, contexts, and movement goals at higher levels that can be flexibly combined with modular ‘motor primitives’ consisting of simple movement patterns at lower levels (*51–53*). Within this perspective, anticipated outcomes may be represented and evaluated before action execution. However, these accounts have typically emphasized hierarchical and compositional descriptions of action control (*9*), without explicitly proposing that goals, action, and anticipated outcomes are integrated non-linearly into conjunctive representations. In contrast, integrative theories explicitly propose that context, goals, actions, and anticipated outcomes are integrated into task-specific event files (*12*, *13*, *15*, *67*). Conjunctive representations such as event files could arise from non-linear mixed selective neuronal populations (*40*, *41*, *68*), enabling a linear readout of multiple task-relevant variables and reducing interference between variables by orthogonalizing their representations (*69–72*). In our results, we find evidence for both compositional and conjunctive representations, suggesting that the brain maintains these formats concurrently. As time progressed, the strength of the compositional stimulus and action representations decreased, whereas conjunctive representations remained relatively stable. This may imply that the brain increasingly relied on task-specific, conjunctive representations to guide action selection (*60*, *64*, *73*).

Surprisingly, the decrease in strength of compositional representations was not coupled with an increase in the strength of conjunctive representations. Moreover, while the strength of action, goal-stimulus-action, and goal-stimulus-action-outcome representations predicted faster RTs on individual trials, the general decrease in RTs as the task progressed could not be explained by an increase in representational strengths. This suggests that the predictive value of the representational structure for behavior does not reflect general practice effects, but other sources of trial-to-trial variability. What, then, could these sources be? Prior work provides a compelling clue: behavioral performance depends on the history of feature overlap across trials with different conjunctions. That is, behavioral interference is greater when some, but not all of the integrated features are repeated from one trial to the next (*15*, *65*). Other work indicates that multiple conjunctive representations can be maintained in parallel, with their strength depending on behavioral relevance (*54*, *74*) or task uncertainty (*63*). Here, we see congruence with influential proposals that action selection reflects a continuous competition between concurrently active sensorimotor patterns rather than a serial evaluation of isolated components (*75*, *76*). Such views of action selection are consistent with event file theory (*12*, *14*) and active inference accounts (*23*, *24*, *77*, *78*), and favor a more integrative perspective on brain function.

Future studies are necessary to determine whether trial-to-trial fluctuations in the strength of conjunctive representations are shaped by additional factors next to trial-history effects, such as uncertainty about action-outcome contingencies or other task demands (*73*, *79*). While our results show a strong relationship between such fluctuations and the fluency of action selection, our method does not permit us to establish a definitive causal link. Other open questions arise from methodological limitations of the current work. One such limitation concerns the arbitrary and variable action-outcome relationships in the experimental task, which may be learned and maintained differently in the brain than more naturalistic and stable action-outcome relationships. Future work should investigate whether the integration of predicted outcomes generalizes to more naturalistic tasks using continuous action spaces, where outcomes unfold dynamically rather than as discrete events. In addition, the current study was limited in its ability to identify the anatomical basis of conjunctive representations due to the relatively poor spatial resolution of EEG measurements. Further research could address this issue by employing large-scale neural recordings to investigate whether distinct cortical or subcortical circuits preferentially support action-outcome integration and whether the dimensionality of neural representations differs across brain areas. For example, a recent fMRI study found that during multi-task learning, conjunctive representations originated in subcortical areas such as the hippocampus and cerebellum, and gradually spread across cortex over the course of learning (*64*). Employing similar approaches in the study of goal-directed action selection will be essential for developing a mechanistic account of how integrated sensorimotor representations support flexible, goal-directed behavior in real-world settings.

Taken together, our findings provide compelling support for the notion that goal-directed action selection operates through the control of predicted sensory outcomes by showing that performance in an action-outcome learning task systematically depends on the trial-to-trial strength of conjunctive neural representations that bind together goals, stimuli, actions, and anticipated sensory outcomes. This dependence underscores the deeply intertwined nature of sensory and motor processes in the brain, and suggests that predicted outcomes are embedded within the very representational architecture that supports action selection.

## Materials and methods

### Participants

40 participants (28 female, 11 male, 1 non-binary; 2 left-handed), aged 18-28 years (*M* = 20.90, *SD* = 2.73), took part in the study. All reported no history of neurological or psychiatric disorders and normal or corrected-to-normal vision. Before participation, all participants provided written informed consent. This study was approved by the Scientific and Ethical Review Board (VCWE) of the Vrije Universiteit Amsterdam. Participants received either financial compensation (10 euros per hour) or course credit.

### Experimental task

Participants performed an action-outcome learning task in which they were asked to generate specific sensory outcomes (Fig 1A). Outcomes were defined by two independent visual features: their shape (circle or star) and spatial frequency (high or low). Low spatial frequency outcomes (1 cycle/degree) were displayed at an orientation of 135 degrees, and high frequency outcomes (4 cycles/degree) at an orientation of 45 degrees, enhancing neural and perceptual discriminability (*80*). All four outcomes subtended 4 × 4 degrees of visual angle at a viewing distance of 70 cm and were matched in terms of overall luminance and surface area. In every block, each unique outcome was mapped to one of four screen locations on a 2 × 2 grid. Participants could generate an outcome by moving a white, square-shaped stimulus from its starting location horizontally (left/right) or vertically (up/down) towards the location of the outcome they wished to generate. Importantly, they first had to learn the state-outcome mapping in a given block, i.e., which combination of stimulus location and action produced which outcome. Due to the organization of the state-outcome mappings (one feature (e.g., shape) varied between top and bottom half of the screen, while the other feature (e.g., spatial frequency) varied between left and right half of the screen), each outcome feature could be reached from each stimulus starting location. Thus, goals, stimuli, and actions were fully orthogonal, and the same goal-stimulus-action combination could occur with two different outcomes. This design enabled the separation of goal-related and outcome-related neural representations.

Each trial started with a goal cue (e.g., ‘circle’) presented for 250 ms, instructing participants which specific outcome feature to generate. Importantly, we used two synonyms to refer to each goal feature (e.g., circle and round) to ensure that goal decoding would not be driven by low-level visual properties of words. After a delay of 250 ms, a white square (‘stimulus’) occurred in one of the four grid locations for 150 ms. The participant then had to generate the outcome with the goal feature by pressing one of two buttons on the keyboard (“X” or “M”), corresponding to a ‘horizontal’ or ‘vertical’ movement (assignment counterbalanced across participants), within 3000 ms from stimulus onset. The chosen action triggered the presentation of the resulting outcome for 250 ms. If no action was selected within the 3000 ms, participants were prompted to respond faster. Each trial ended with a jittered interval of 300 to 500 ms.

The experiment consisted of two sessions, each starting with a brief familiarization with the goal cues, followed by two practice blocks of at least 64 trials and 12 experimental blocks of at least 160 trials. Practice blocks followed a slower pace and contained feedback on every trial. Within the 12 experimental blocks, we used four different state-outcome mappings that each occurred in three non-consecutive blocks. These four mappings were chosen so that across the experiment, each outcome occurred equally often in each of the four locations, preventing decoding biased by associations between specific outcomes and locations. Blocks were organized pseudo-randomly such that the first 80 trials and second 80 trials would each contain all 16 possible goal-stimulus-action-outcome combinations exactly five times, in shuffled order. This was done to ensure that the first and second half of each experimental block would contain the same number of trials from each trial type, facilitating comparison. Each block continued until at least 160 trials had been completed and accuracy exceeded 85% on the last 16 trials, ensuring that participants had fully learned the state-outcome mapping by the end of the block.

### EEG recording and preprocessing

Continuous EEG was recorded using a Biosemi ActiveThree system with 64 electrodes arranged according to the international 10-20 system. Four EOG electrodes were added to monitor vertical and horizontal eye movements and two additional electrodes were placed on the left and right mastoids. Data was digitized at 512 Hz and down-sampled offline to 128 Hz. For preprocessing, data was re-referenced to the average of the mastoids, high-pass filtered at 0.01 Hz, and eyeblink artifacts were removed using ICA. We then extracted wide trial-based epochs from -300 ms to 6000 ms relative to goal cue onset, using the period between -200 ms and -50 ms for baseline correction. From these broad epochs, we extracted stimulus-locked (- 600 ms to 800 ms relative to stimulus onset) and action-locked (-800 ms to 600 ms relative to action) epochs. All EEG data was processed using MATLAB-based software EEGLAB (*81*)

### Representational similarity analysis

The experimental design orthogonalized task features as much as possible, resulting in 64 unique trial combinations (4 mappings × 4 goal cues × 4 stimulus locations). However, as stimuli could be moved horizontally or vertically, but not diagonally, stimulus location and outcome location were not fully independent (e.g., if the stimulus was located in the upper-right location, the outcome could only occur in the lower-right or upper-left location, but not the upper-right or lower-left location). To fully orthogonalize stimuli and outcomes, trials were divided into two sets: one containing all 32 trial combinations with the stimulus in the upper-left or lower-right location and one containing all 32 trial combinations with the stimulus in the upper-right or lower-left location (Fig 1B). Within each set, all task features were fully independent, and analyses were therefore performed separately for each set.

Following Kikumoto and Mayr (2020), we performed time-resolved multivariate pattern analysis using the ADAM toolbox (*82*) on EEG data from correct trials only. For each participant and each timepoint, a backward decoding model was trained on raw EEG data, using 5-fold cross-validation in which each fold served as the testing set once. The number of observations from each trial combination (i.e., combination of goal, stimulus location, action, outcome identity, and outcome location) was equated within training sets by randomly dropping excess trials. The entire cross-validation procedure was repeated eight times, newly generating five randomized folds on each iteration. We defined the final model classification scores as the average trial- and time-specific classification scores over all eight iterations.

Next, we applied representational similarity analysis (RSA; Fig 1C) on the model classification scores to track the underlying similarity structure of task features and their combinations over time. We constructed model RSA matrices for each individual task feature (goal, stimulus location, action, outcome identity) as well as for the combinations of goal-stimulus-action and goal-stimulus-action-outcome. We included additional RSA model matrices for location-outcome mappings and for outcome locations, and a vector of subject-specific mean RTs for each unique trial combination as nuisance regressors. For each trial and time sample, we regressed the log-transformed model classification scores for each possible trial combination onto the RSA model vectors for the true trial combination label and nuisance regressors, entering all predictors simultaneously to isolate their unique contributions. The resulting *t*-values served as a measure of representational pattern strength. For each time sample, we excluded *t*-values that were more than ±5 SDs away from the mean, resulting in the exclusion of 0.04% of all estimated values.

### Linear mixed-effects modeling and cluster-based permutation testing

To assess the statistical robustness of our findings, we combined linear mixed-effects modeling with nonparametric cluster-based permutation testing (cf. Maris & Oostenveld, 2007). We fitted linear mixed-effects models to quantify behavioral learning effects across sessions, blocks, and trials, and to relate RSA-derived coefficients to trial-to-trial variability in RTs. We further fitted linear models to quantify learning effects in RSA-derived coefficients, both within and across blocks. Prior to model fitting, trials with RTs that were implausibly fast (< 100 ms) or excessively slow (> 5 SDs from grand mean) were removed, resulting in the exclusion of 0.17% of all trials. All models were fitted using a subject-specific intercept as a random effect. For models with RT as the dependent variable, RTs were log-transformed to reduce skew. For models with accuracy as the dependent variable, we fitted logistic mixed-effects models using a logit link-function. To establish learning, we inspected both the main effects and interaction effects of sessions, blocks, and trials. Finally, to quantify learning effects in RSA-derived coefficient values, we fitted linear models using RSA-coefficients as dependent variables and log-transformed trial position as the predictor (counted from the start of the current block for within-block learning and from the start of the experiment for across-block learning). All models were constructed and fitted using the R-based package *lme4* (*83*).

For time-resolved effects, we used nonparametric cluster-based permutation testing (*84*) to identify significant clusters in RSA time courses, in their relationship with trial-to-trial RTs, and in their development across trials within single blocks. Empirical clusters were defined as seven or more consecutive time points (>50 ms) for which *t*-values exceeded the cluster-forming threshold of *p* < 0.05, and cluster-level statistics were computed by summing *t*-values within each cluster. Then, we constructed a nonparametric null distribution by computing cluster-level statistics across 1000 permutations with shuffled condition labels. Empirical clusters were considered significant if their cluster-level statistic exceeded the 99th percentile of the null distribution (*p* < 0.01). For analyses of RSA scores across time and their relationship to trial-to-trial RTs, each permutation involved repeating the full multivariate pattern analysis on EEG data with shuffled trial combination labels, followed by RSA and linear mixed modelling on the resulting model classification scores. For the assessment of RSA changes across trials, permutations consisted of refitting the linear mixed model with shuffled trial-position labels.

## Acknowledgements

We are grateful for the assistance of A. Cavalli during data collection.

## Data, code, and materials availability

Data and analysis codes are available at https://doi.org/10.17605/OSF.IO/4PWK7.

## Supplementary Materials for

**Fig S1.**
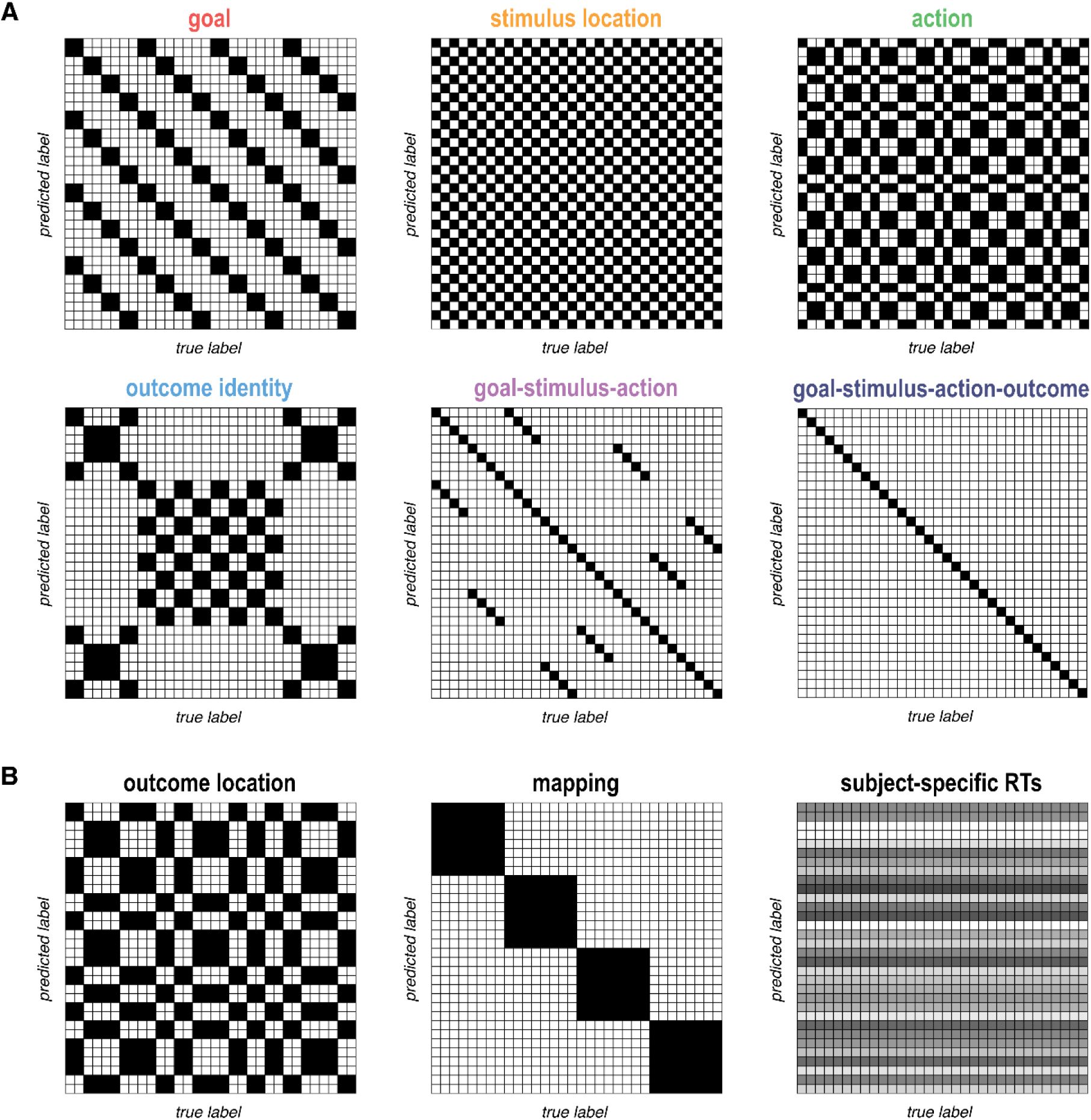
RSA models. **A** Full model RSA matrices for predictors of interest: goal, stimulus location, action, outcome identity, goal-stimulus-action conjunctions, and goal-stimulus-action-outcome conjunctions. The two conjunctive predictors code for non-linear interactions between features that are not captured by the linear addition of single features. **B** Full model RSA matrices for control variables: outcome location, action-outcome mapping, and normalized subject-specific reaction times for each trial combination (showing values of one example participant). Note that all matrices are 32 × 32, representing the 32 unique trial combinations in each set.

**Fig S2.**
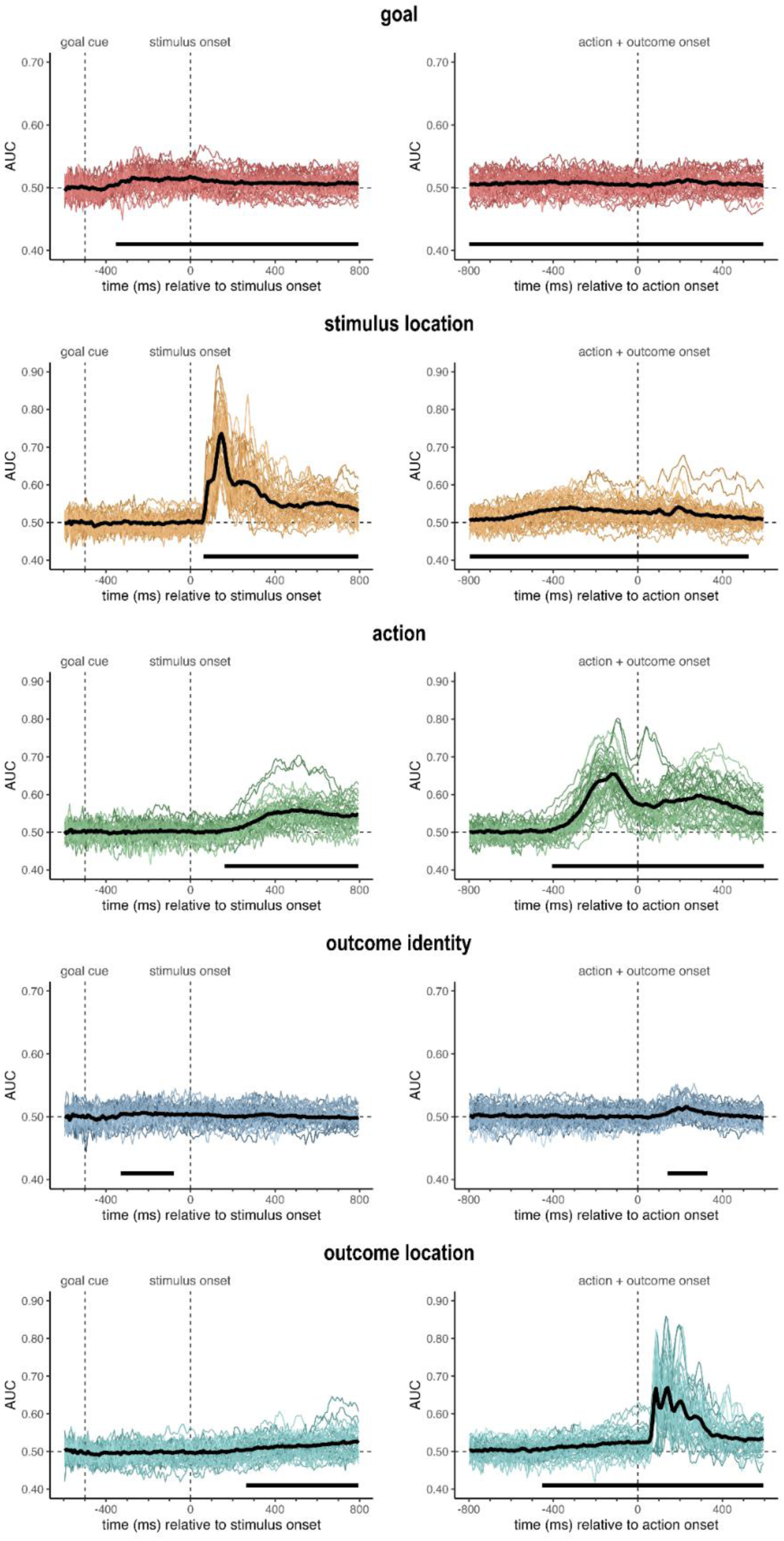
Feature decoding results. Cross-validated classifier decoding results for single features, separately for stimulus-locked and action-locked epochs. Black lines represent grand average, colored lines represent individual participants. Black bars beneath each panel denote significant clusters identified via nonparametric cluster-based permutation testing (10,000 permutations; cluster-forming threshold: *p* < 0.05; cluster significance threshold: *p* < 0.05). Note that the maximum Y-axis value may differ across features (0.70 or 0.90).

**Fig S3.**
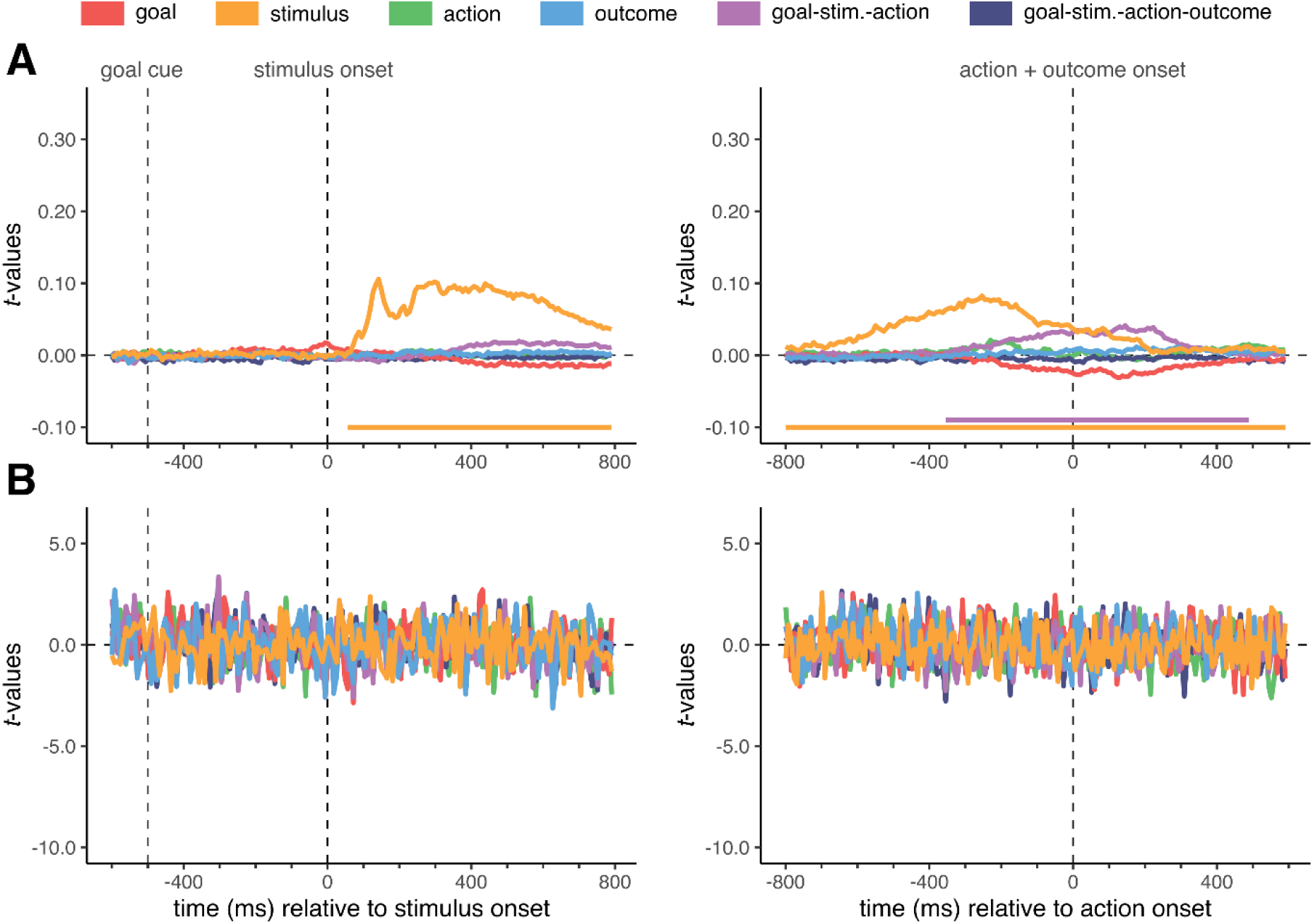
Representational EOG time course. **A** Average trajectory of single-trial *t*-values derived from representational similarity analysis (RSA) using EOG and eye tracking channels, time-locked to stimulus onset (*left*) and action onset (*right*). **B** Trajectory of *t*-values from mixed linear models predicting trial-to-trial RTs from eye-RSA scores, time-locked to stimulus onset (*left*) and action onset (*right*). Colored bars at the bottom of each panel indicate significant clusters, derived from nonparametric cluster-based permutation testing (1000 permutations; cluster-forming threshold: *p* < 0.05; cluster significance threshold: *p* < 0.01).

